# Whole-Genome Sequencing of Three Extremophile *Bacillus sp*. Strains Isolated from the Atacama Desert

**DOI:** 10.1101/2024.02.11.578748

**Authors:** Nicole T Cavanaugh, Girish Kumar, Matteo Couto Frignani, Nathan Thewedros, Marcello Twahirwa, Carlos Riquelme, André O Hudson, Yunrong Chai, Veronica Godoy-Carter

## Abstract

The Atacama Desert is home to bacteria that use biofilms as a means of protecting themselves against the harsh environment. We cultured and sequenced the genomes of three *Bacillus sp*. isolates from the soil of the Atacama Desert. This information will inform research in survival mechanisms of eubacteria in the Atacama Desert.

## Announcement

The Atacama Desert is one of the most ancient, most highly irradiated, and driest deserts in the world (1). Despite this, it is home to a variety of bacterial life. Bacteria in this region are shown to have developed adaptations to survive this harsh, irradiated environment such as high levels of pigment production and biofilm formation (2). In the past, most research addressing survival mechanisms in microbes from the Atacama have focused on archaea. We are studying how eubacteria have developed adaptations to survive in such a desert environment.

To study how biofilms impact bacterial survival, we started by culturing and further isolating a variety of bacterial colonies from the desert soil and screening for biofilm-forming isolates. Bacterial colonies were purified after plating 100 μL of a soil water slurry on Reasoner’s 2A (R2A) agar medium and incubating plates at 28 ºC for 48-72 hours. To identify the isolates, we performed 16S rRNA gene sequencing on the V3/4 variable regions using the following primers: 5’-CCTACGGGNGGCWGCAG-3’ and 5’-GACTACHVGGGTATCTAATCC-3’. Using National Society for Biotechnology Information (NCBI) BLASTN version 2.15.0, three *Bacillus sp*. were identified (3, 4). The *Bacillus* genus contains Gram-positive bacteria that are found throughout the environment, many of which are known to produce biofilms (5, 6, 7, 8, 9, 10). To test biofilm formation, strains were grown at 37 ºC to OD600 = 1.0 in Luria-Bertani (LB) broth and 2 μl of culture was spotted on LB agar plates supplemented with 1% glycerol and 0.1 mM MnSO_4_ (LBGM, biofilm inducing medium) (11). Colony biofilms were incubated for 48 hours.

**Figure 1:**
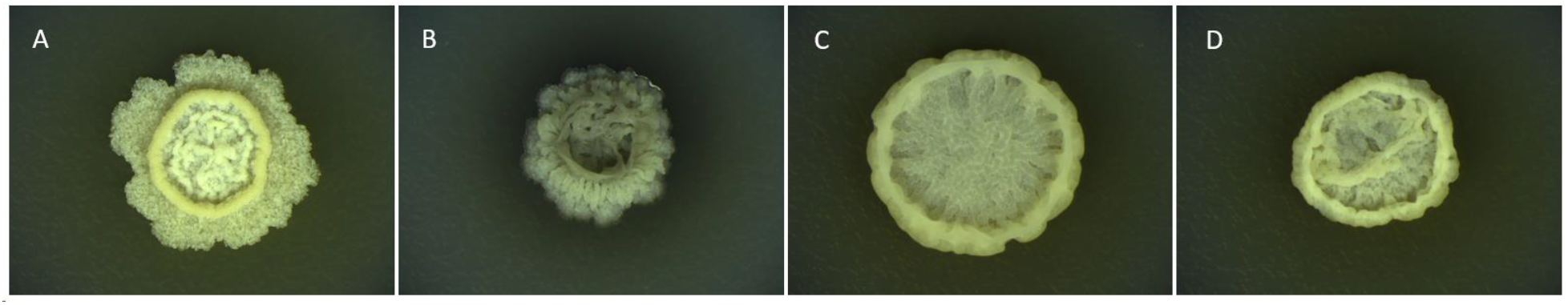
Environmental strains isolated from the Atacama Desert form colony biofilms. Colony biofilms of 0102A (A), 0209A (B), 0909A (C), and 3610 (B. subtilis lab strain) (D) grown on LBGM medium (11) after 48 hours of incubation.

These *Bacillus sp*. were selected for further genomic analysis. Genomic DNA was isolated from a 2-mL LB culture using the Wizard Genomic DNA Purification Kit (Promega, USA) following kit instructions for Gram-positive bacteria. Purified DNA was quantified using a NanoDrop spectrophotometer (Thermo-Fisher), DNA libraries were prepared using the Nextera XT library preparation kit (Illumina) and sequenced using the Illumina MiSeq System at the Genomics Lab, Rochester Institute of Technology. Raw reads were trimmed using Trimmomatic Galaxy version 0.38.1 (12). The trimmed reads were assembled de novo using Unicycler Galaxy version 0.5.0+galaxy1 (13). Quality analysis was performed using Quast Galaxy version 5.2.0+galaxy1 (14) and FastQC Galaxy version 0.74+galaxy0 (15). Quality assessments are outlined in Table 1. All programs used were run at default parameters unless otherwise noted. Analysis was performed using the Prokaryotic Genome Analysis Pipeline (PGAP) version 6.6 to reveal the total number of genes, protein coding sequences, rRNAs, and tRNAs in each assembly (Table 1) (16, 17, 18). Further analysis of the *eps* operon in each genome using NCBI BLASTN Galaxy Version 2.14.1+galaxy and taxonomy analysis by the Automated Multi Locus Species Tree (autoMLST) showed that these three isolates were most closely related to *Bacillus vallismortis and Bacillus mojavensis* (3-4, 19).

**Table 1:** Summary of quality assessments for each strain. A summary of the identities and quality assessments of each strain 0102A, 0209A, and 0909A. For each strain, the total bases and number of reads was determined using FastQC Galaxy version 0.74+galaxy0. Read length, number of contigs, N50, and GC content were assessed using Quast Galaxy version 5.2.0+galaxy1. Numbers of total genes, protein coding sequences, rRNAs, and tRNAs were determined using the Prokaryotic Genome Analysis Pipeline (PGAP) version 6.6.

| Isolate | Most Closely Related Species | Total Bases (Mbp) | Read Length (bp) | Number of Reads (FastQC) | Number of Contigs | N50 (bp) | GC Content | Total Genes | Protein Coding Sequences | rRNA Operons | tRNAs |
| --- | --- | --- | --- | --- | --- | --- | --- | --- | --- | --- | --- |
| 0102A | *B. vallismortis* | 813.2 | 4,192,438 | 4,406,089 | 27 | 350,970 | 43.87% | 4296 | 4067 | 3 | 76 |
| 0209A | *B. vallismortis* | 980.4 | 4,167,904 | 5,239,849 | 19 | 1,048,338 | 43.98% | 4208 | 3989 | 3 | 73 |
| 0909A | *B. mojavensis* | 857.1 | 3,899,559 | 4,586,376 | 19 | 930,776 | 43.66% | 4061 | 3870 | 3 | 67 |

## Data Availability

The WGS projects for *Bacillus subtilis* strains 0102A, 0209A, and 0909A have been deposited in GenBank under BioProject number PRJNA1071648. Strain 0102A is deposited under BioSample number SAMN39706723 and SRA number [SRR27839954](https://www.ncbi.nlm.nih.gov/sra/SRX23503239%5baccn%5d). PGAP analysis can be found under accession number JBAIZW000000000. Strain 0209A is deposited under BioSample number SAMN39706724 and SRA number [SRR27839953](https://www.ncbi.nlm.nih.gov/sra/SRX23503240%5baccn%5d). PGAP analysis can be found under accession number JBAIZV000000000. Strain 0909A is deposited under BioSample number SAMN39706725 and SRA number [SRR27839952](https://www.ncbi.nlm.nih.gov/sra/SRX23503241%5baccn%5d). PGAP analysis can be found under accession number JBAIZU000000000.

The [NCBI BLASTN](https://blast.ncbi.nlm.nih.gov/Blast.cgi?PROGRAM=blastn&BLAST_SPEC=GeoBlast&PAGE_TYPE=BlastSearch) webserver was used to identify the 0102A, 0209A, and 0909A 16s rRNA gene sequence. All programs found on the Galaxy platform can be found at usegalaxy.org.

## Acknowledgements

We would like to thank the National Science Foundation (MCB1651732) and the NU Global Experience Office for their support. Thank you to the RIT Genomics Lab. V.G. was supported by the HHMI Inclusive Excellence grant from the NU Biology Department. M.C.F, N.T., and M.T. were supported by NU Undergraduate Research and Fellowships Office PEAK Awards. N.C. is supported by the NSF Graduate Research Fellowship Program (1938052).

